# Alk1/Endoglin dependent increases in vein endothelial cell sizes precipitate arteriovenous malformations

**DOI:** 10.1101/2023.11.07.566079

**Authors:** Zeenat Diwan, Arndt F. Siekmann

## Abstract

Aberrations in blood vessel diameters can disrupt the hierarchical patterning of the vasculature and cause congenital vascular anomalies, such as arteriovenous malformations (AVMs). Despite the identification of the Bone Morphogenetic Protein (BMP) pathway as a major driver in AVM pathology, we still lack an understanding of the early embryonic events regulating vessel hierarchy and arteriovenous shunt formation *in vivo*. We therefore studied blood vessel diameter control of the dorsal aorta (DA) and posterior cardinal vein (PCV) in zebrafish embryos. Our findings reveal that increases in blood flow during embryonic development result in increases in arterial endothelial cell (EC) sizes, ultimately enlarging DA diameters. By contrast, anterior regions of the PCV did not respond to changes in blood flow, while caudal regions, close to the artery-vein junction, responded to changes in flow like the DA, but to a lesser extent. To unravel the mechanisms underlying the reduced response of PCV ECs to flow, we studied zebrafish embryos mutant for the BMP pathway components *endoglin* and *alk1*. Through the generation of genetic mosaics, we discovered that both Endoglin and Alk1 were required cell autonomously in PCV cells to restrict EC sizes and thereby limit venous diameter increases in response to flow. We further revealed that initial increases in the diameter of the caudal PCV secondarily led to increased DA diameters and cell sizes. Therefore, Alk1/Endoglin signaling prevents vein ECs from behaving like arterial ECs. This differential response of arterial and venous EC cells to increases in flow is necessary to prevent the development of AVMs. This study thus offers insights into the spatiotemporal regulation of vessel hierarchy during early development and identifies changes in EC shapes as an important contributor in determining blood vessel diameters. Failure in this mechanism underlies the vein-specific initiation of AVMs in vertebrate models of HHT.

## Introduction

Biological tubes are the fundamental structural units of many organs, including the vascular system. A hierarchical organization of correctly sized blood vessels ensures an appropriate distribution of blood to all parts of the body. Aberrations in the diameter of these vessels can disrupt this hierarchy and cause congenital vascular disorders, including Hereditary Hemorrhagic Telangiectasia (HHT), Cerebral Cavernous Malformations (CCM) and Venous and Lymphatic Malformations (Fernandez-Flores et al., 2023). HHT is characterized by abnormal arterial and venous vessel dilations that cause direct connections between the vessels and precipitate as shunts, called arteriovenous malformations or AVMs. These AVMs often reduce the quality of life of the individual or at times may prove fatal (Sobrepera et al., 2021). The underlying molecular mechanisms involve a loss of function of BMP pathway components, such as Endoglin, Alk1, Smad4 or BMP9 (Gallione et al., 2004; Johnson et al., 1996; McAllister et al., 1994; Wooderchak-Donahue et al., 2013). However, despite advances in elucidating the genetic and molecular basis underlying AVMs, a detailed *in vivo* understanding of the spatiotemporal mechanisms that regulate vessel diameters during development and how their disruption initiates AVM formation is still lacking.

Previous research has noted that alterations in the magnitude of flow, and consequently fluid shear stress (FSS), result in corresponding modifications in the diameter of arteries. This phenomenon has been suggested as the shear stress set point theory, which proposes that FSS below or above a specific threshold value initiates vessel remodelling: constricting for low FSS and dilating for high FSS, to return to the original shear stress level and uphold homeostasis. However, the existence of a shear stress set point for both arteries and veins would mean that the unchecked increase in blood flow during embryonic development would result in continual increases in vessel diameters of the shortest connection between an artery and a vein, leading to arteriovenous shunt formation (Pries et al., 2010; Thoma, 1894). Since this does not happen during normal development, mechanisms must exist to counteract the effects of shear stress set point-driven blood vessel dilations or mechanisms that impart this set point must exist only for selective vessel types. Interestingly, although the set point theory is widely recognized as applicable to all vessels, an *in vivo* shear stress set point has so far only been shown for arteries (Baeyens et al., 2015; Langille, 1996; Tuttle et al., 2001). Whether an *in vivo* set point similarly exists for veins remains to be explored.

Several *in vitro* and a few *in vivo* studies over the past years have revealed that most types of vascular malformations, including AVMs, have a key pathological characteristic in common – the aberrant vessel dilations result from an increase in endothelial cell (EC) number or size or both. A considerable focus has been put on EC number (through proliferation or migration) as being the prominent effector of vessel diameter changes during pathogenesis (Castillo et al., 2016; Crist et al., 2018; Jin et al., 2017; Li et al., 2018; Poduri et al., 2017; Rochon et al., 2016; Tual-Chalot et al., 2014); however we and others have shown that EC sizes play an equally important – if not a bigger – role (Crist et al., 2018; Murphy et al., 2014; Poduri et al., 2017; Sugden et al., 2017). Despite this knowledge, we still lack sufficient insight into the *in vivo* mechanisms through which EC size regulates vessel diameter in a spatiotemporal manner and how failure of this regulation might contribute to the pathogenesis of AVMs.

To investigate these open questions, we performed a detailed analysis of changes in vessel diameters and EC morphologies during zebrafish embryonic development. Intriguingly, we discovered that while the artery consistently responds to changes in blood flow and is dependent on flow increase for expansion of its diameter and EC sizes, the vein shows context-dependent responses. The anterior part of the vein closest to the hart lacks observable changes, while the region close to the artery-vein junction responds to an increase or decrease in flow like the artery, but to a lesser extent. Our data thus support the shear stress set point theory for arteries, but curiously also implicate that an *in vivo* set point for veins might be spatially regulated during early development.

To ascertain the molecular mechanisms that regulate venous growth (or allow for arterial growth) in response to flow, we investigated the BMP receptor Alk1 and the co-receptor Endoglin. We find that both Endoglin and Alk1 are required for EC size control specifically in the vein. We also reveal that in *endoglin* (*eng*) mutants, vessel dilation initiates in the major vein and secondarily causes expansion of the artery, leading to shunt formation. Overall, our data support a model wherein activation of Endoglin and Alk1 signalling makes veins less responsive to flow and thus limits venous diameter increase to prevent shunt formation during early development. Our findings uncover a previously under-appreciated role for vessel-specific regulation of diameters and EC sizes during early development and shed light on the cellular mechanisms underlying the spatiotemporal initiation of AVMs in a vertebrate model for HHT.

## Results

### The dorsal aorta and posterior cardinal vein show differential growth during early development

To investigate whether an *in vivo* shear stress set point exists for both arteries and veins, we examined changes in vessel diameters in response to variations in flow magnitude during early zebrafish development. We first performed timelapse imaging of the major artery (dorsal aorta, DA) and major vein (posterior cardinal vein, PCV) in the zebrafish embryo starting at 1 day post fertilization (dpf; when the vessels have just formed and lumenized) through 3dpf (when vessel remodeling occurs)). We focused on two adjacent vessel beds in the axial vasculature – the anterior trunk and posterior tail regions **(Fig. 1A, B)**. We observed that in the trunk region, the vein started out with a significantly larger diameter than the artery at 1dpf (p<0.0001) **(Fig. 1C)**. More importantly, while the artery showed a dramatic increase in diameter from 1-2 dpf in correlation with increase in blood flow magnitude (Santoso et al., 2019), the vein showed minimal fluctuations during this time interval. To gain deeper insights into the EC parameters that effect change in arterial and venous diameters, we performed a detailed morphometric analysis of both trunk and tail arterial and venous ECs. To achieve this, we utilized a previous strategy wherein we imaged *Tg(fli1a:lifeactGFP)^mu240^*embryos that have EC junctions marked with EGFP (Sugden et al., 2017). This allows us to trace the boundaries of individual ECs and quantify their morphologies. Interestingly, changes in EC sizes for both the artery and vein were concurrent with diameter changes from 1-2 dpf **(Fig. 1D)** and indicated that expansion of vessel diameter during early development occurs through an increase in EC size. No significant changes in EC size were noticed between 2 and 3 dpf. However, during this period, arterial and venous ECs aligned to the vessel axis **(Suppl. Fig. 1A-D)**, correlating with a decrease in vessel diameter **(Fig. 1C)**.

**Figure 1.**
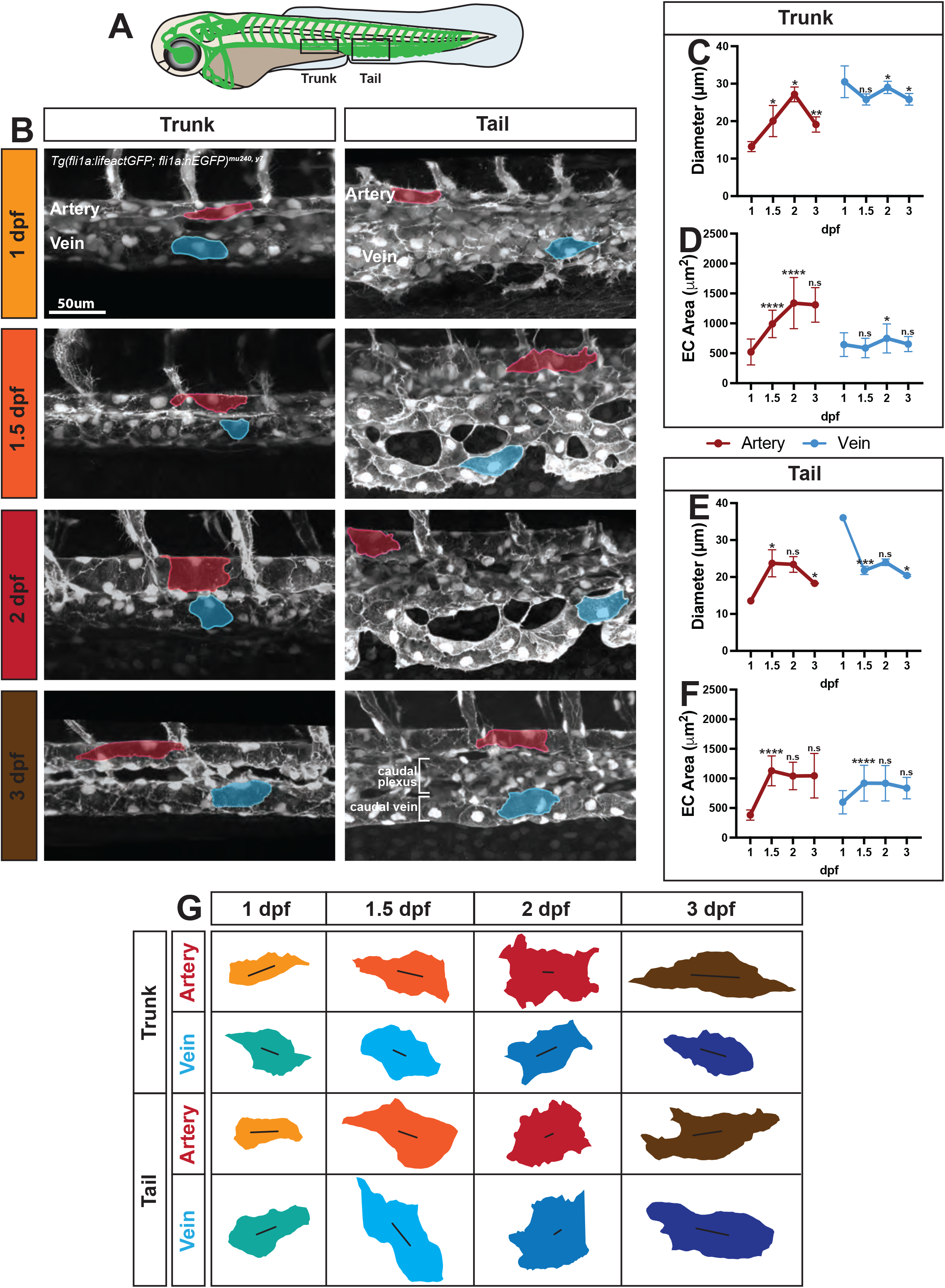
The major artery and vein show differential growth during development. (A) Schematic of a 3dpf WT zebrafish embryo with the trunk and tail regions outlined by solid black boxes. (B) Timelapse images of the major artery and vein in the trunk and tail regions of WT embryos at 1, 1.5, 2 and 3dpf. Representative ECs outlined to show size changes. Red = arterial EC, blue = venous EC. (C-F) Quantifications of vessel diameter and EC area of the major artery and vein in the trunk (C, D) and tail (E, F) regions. Data analyzed across developmental stages by two-way ANOVA. n.s, not significant; *P <0.05, **P <0.01, ***P <0.001, ****P <0.0001; error bars indicate s.d. (G) 2D cell outlines of arterial and venous ECs at 1, 1.5, 2 and 3dpf in the trunk and tail regions. Length and angle of black line within each cell represents elongation and alignment of the cell, respectively.

To explore the contribution of EC number to changes in vessel diameter, we quantified EC nuclei in the trunk region using *Tg(fli1a:nEGFP)^y7^* embryos. We discovered that both arterial and venous EC numbers (measured per 430 um length of vessel) showed a significant decrease over time (1-3dpf) **(Suppl. Fig. 1E, G)** correlating with migration of ECs out of the vessels to contribute to other cell types like arterial or venous ISVs, lymphatics and hematopoietic cells (Eberlein et al., 2021; Isogai et al., 2003; Rosa et al., 2022). Importantly, the vein had significantly higher EC numbers compared to the artery at all four developmental stages analyzed between 1-3 dpf. Additionally, trunk venous ECs showed around 15-30% proliferation rate during this time **(Suppl. Fig. 1F, H, I, J)**. Lastly, although we observed higher proliferation events in the trunk vein than in the artery, venous EC proliferation rate (number of proliferation events/number of ECs) was not significantly different from that of the artery between 1-3 dpf **(Suppl. Fig. 1F)**. This suggests that EC proliferation is unlikely to influence diameter differences between the two vessels. In conclusion, our observations indicate that the trunk artery grows in diameter through an increase in EC size. On the other hand, the trunk vein starts out with a larger diameter – a consequence of higher EC number – and undergoes marginal changes over time.

Like the trunk vessels, the tail artery and vein started out with similar EC sizes, and with arterial ECs showing a significantly higher growth rate than venous ECs (2-fold growth in arterial EC size versus 0.5-fold growth in venous from 1-1.5 dpf) **(Fig. 1F)**. The initial increase in arterial EC size correlated with an expansion of arterial diameter **(Fig. 1E)**. Also like the trunk, the tail vein started out significantly larger than the artery at 1 dpf – a consequence of higher EC numbers. However, unlike the trunk vein that is formed by vasculogenesis and remains relatively unchanged over time, the tail vein starts out as an ovoid structure that undergoes extensive angiogenic remodelling between 1-2 dpf to give rise to two distinct but connected structures – the caudal plexus and the caudal vein **(Fig. 1B, see 3 dpf tail image)**. The caudal vein, which starts to form around 1.5 dpf, had a significantly smaller luminal diameter than the ovoid vein structure at 1dpf **(Fig. 1E**). Moreover, it behaved similarly to the trunk vein – there were minimal changes in its diameter and EC size and a decrease in EC number over time **(Fig. 1E, F; Suppl. Fig. 1G**). Lastly, EC proliferation rate was not significantly different between the tail artery and vein **(Suppl. Fig. 1H)**, which paralleled our observations in the trunk. To summarize, we observe that tail vessels behave very similarly to trunk vessels, in that the artery increases in diameter through an expansion of EC size, while the vein undergoes comparatively smaller changes in its diameter and EC size over time (except for diameter changes during tail vein remodelling between 1-1.5 dpf). Interestingly, the tail artery and vein are more similar to each other than the trunk vessels. Altogether, our findings reveal the existence of spatiotemporal differences in EC morphology in the artery and vein during zebrafish development (**Fig. 1G**). More importantly, while the artery experiences a substantial change – both in diameter and EC size – in correlation with increase in blood flow during early development, the response of the vein to increasing flow is marginal. This suggests that, unlike the artery, the vein might not have a shear stress set point.

### Reducing blood flow during early development affects arterial growth, while venous ECs show differential spatial responses

To further explore our hypothesis that an *in vivo* shear stress set point does not exist for the vein, we decided to study the effect of reducing blood flow on arterial and venous diameters and EC sizes. Treatment of 2 dpf embryos with 500 μM (0.13mg/mL) tricaine can result in a 40% reduction in blood flow velocity (Malone et al., 2007). To achieve a more significant reduction in flow in our study, we thus treated zebrafish embryos with a higher dose of tricaine, 0.5mg/ml, for two hours at each of four developmental stages – 1, 1.5, 2 and 3 dpf. This was followed by imaging to determine vessel diameters and EC sizes in the trunk and tail regions. We observed that in tricaine-treated embryos, trunk arterial ECs showed a significant reduction in their size in correlation with a decrease in vessel diameter at 26, 38 and 50 hours post fertilization (hpf). Conversely, venous ECs were unaffected by reduction in flow, with fairly small changes in venous diameter **(Fig. 2A, B, D)**. The tail artery of tricaine-treated embryos showed similar trends in EC size and vessel diameter as observed in the trunk. However, the tail vein diameter and EC size at 50 hpf were significantly smaller in tricaine-treated embryos compared to control embryos **(Fig. 2C, E)**, indicating that tail vein growth requires blood flow. This contrasted with the trunk vein, which did not seem to be affected by changes in flow. There are two interesting points to note here: first, tail vein diameters of tricaine-treated embryos were significantly larger than those of controls at 26 and 38 hpf **(Fig. 2C, E)**. This was due to inhibition of vessel remodeling in the presence of low or no flow. As noted earlier, the tail vein undergoes extensive remodeling between 24-48 hpf that involves conversion of an ovoid structure into the caudal venous plexus and caudal vein. This remodeling requires blood flow (Goetz et al., 2014; Karthik et al., 2018; Kugler et al., 2021). Thus, experimentally reducing flow during these early development time points causes a failure to remodel, resulting in a single venous structure with a large lumen. Secondly, we noticed that reducing flow after the remodeling period, around 50 hpf, causes a redirection of majority of blood flow into a more dorsal venous channel (outlined by dashed lines in **Fig 2C**) with some flow through the caudal vein. To summarize, our results show that the artery responds to changes in blood flow in both trunk and tail regions in a manner that is consistent with the presence of an *in vivo* shear stress set point during early development. On the other hand, the vein’s response is context-dependent – the trunk vein is not affected by either increases or decreases in flow, and thus does not seem to have a shear stress set point, while the tail vein needs flow for growth and might have an *in vivo* set point like the artery.

**Figure. 2.**
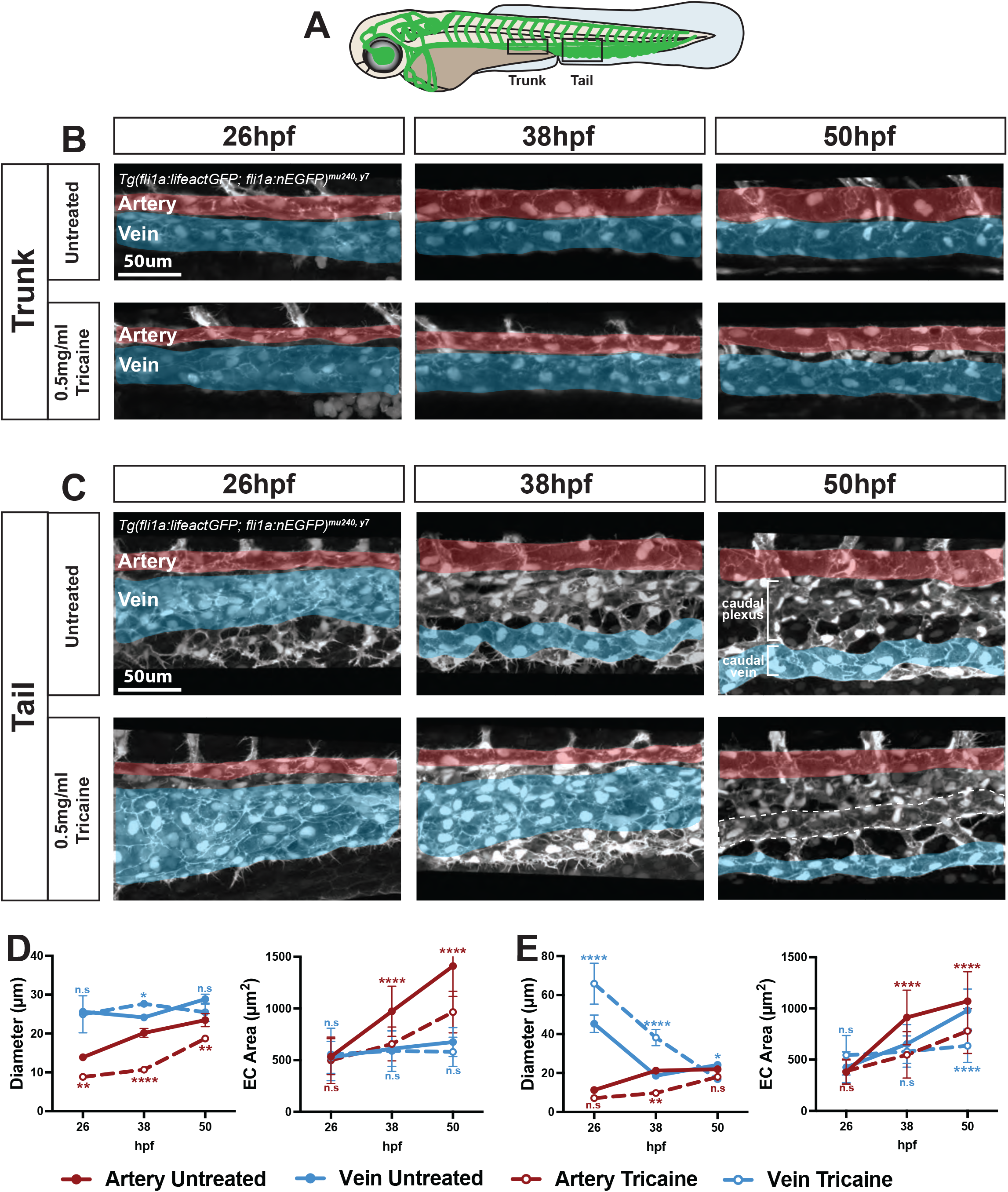
Reducing blood flow affects arterial EC growth, while venous ECs show differential effects. (A) Schematic of a 3dpf WT zebrafish embryo with the trunk and tail regions outlined by solid black boxes. (B,C) Confocal images of the major artery and vein in the trunk (B) and tail (C) regions of WT embryos at 26, 38 and 50hpf either untreated or treated with 0.5mg/ml tricaine for 2h. (D,E) Quantifications of vessel diameter and EC area of the major artery and vein in the trunk (D) and tail (E) regions. Data analyzed across treatment conditions by two-way ANOVA. n.s, not significant; *P <0.05, **P <0.01, ***P <0.001, ****P <0.0001; error bars indicate s.d.

### Global loss of endoglin increases EC size in both the artery and vein

The finding that changes in blood flow resulted in the remodeling of arterial vessels while veins responded in a region-specific manner suggests that there are mechanisms operating in the vein that spatially restrict EC growth in response to flow. To unravel these mechanisms, we investigated the BMP signaling components Endoglin and Alk1. Loss-of-function of either leads to arteriovenous malformations (AVMs) in humans, mice and zebrafish (Baeyens et al., 2016; Jin et al., 2017; Rochon et al., 2016; Sobrepera et al., 2021; Sugden et al., 2017). Studies in mice and zebrafish have furthermore shown that Endoglin is predominantly expressed in veins and capillaries (Jin et al., 2017; Mahmoud et al., 2010; Sugden et al., 2017), while Alk1 is more highly expressed in arteries (Roman et al., 2002; Seki et al., 2003). We thus hypothesized that Endoglin in the vein and Alk1 in the artery might regulate EC size in these compartments in response to flow. We first performed whole mount *in situ* hybridization to confirm the expression pattern of these molecules in the trunk and tail vasculature of zebrafish embryos. As expected, we discovered that Endoglin expression in zebrafish axial vasculature was predominantly localized to the vein, with higher expression in the tail compared to the trunk. Conversely, Alk1 was expressed largely in the artery **(Suppl. Fig. 2A)**.

We previously showed that Endoglin regulates tail artery diameters during zebrafish development by controlling EC shape and size (Sugden et al., 2017). As Endoglin is principally expressed in the vein, we extended our analysis to venous regions of the mutant vasculature **(Fig. 3A, B)**. No significant differences were found in the number, elongation or alignment of *eng* mutant ECs in the trunk **(Suppl. Fig. 3A-C)** or tail vessels **(Suppl. Fig. 3D-F)**. Interestingly, an increase in EC size was observed in both vessel types – specifically the trunk artery **(Fig. 3D)**, tail artery and tail vein **(Fig. 3F)** and correlated with an increase in the diameters of the respective vessels **(Fig. 3C, E)**. We observed a similar percentage of increase (17%) in both trunk and tail arterial ECs in *eng* mutants compared to siblings. However, loss of *eng* differentially affected the trunk and tail vein. While mutant ECs in the trunk vein did not show significant changes, tail vein ECs were 50% larger in mutants compared to siblings. Collectively, these observations indicate that although global loss of Endoglin affects both arterial and venous diameters mainly through an increase in EC size, the most prominent effect is in the tail vein **(Suppl. Fig. 3G, H)** and corresponds with the high expression of Endoglin in this vessel.

**Figure 3.**
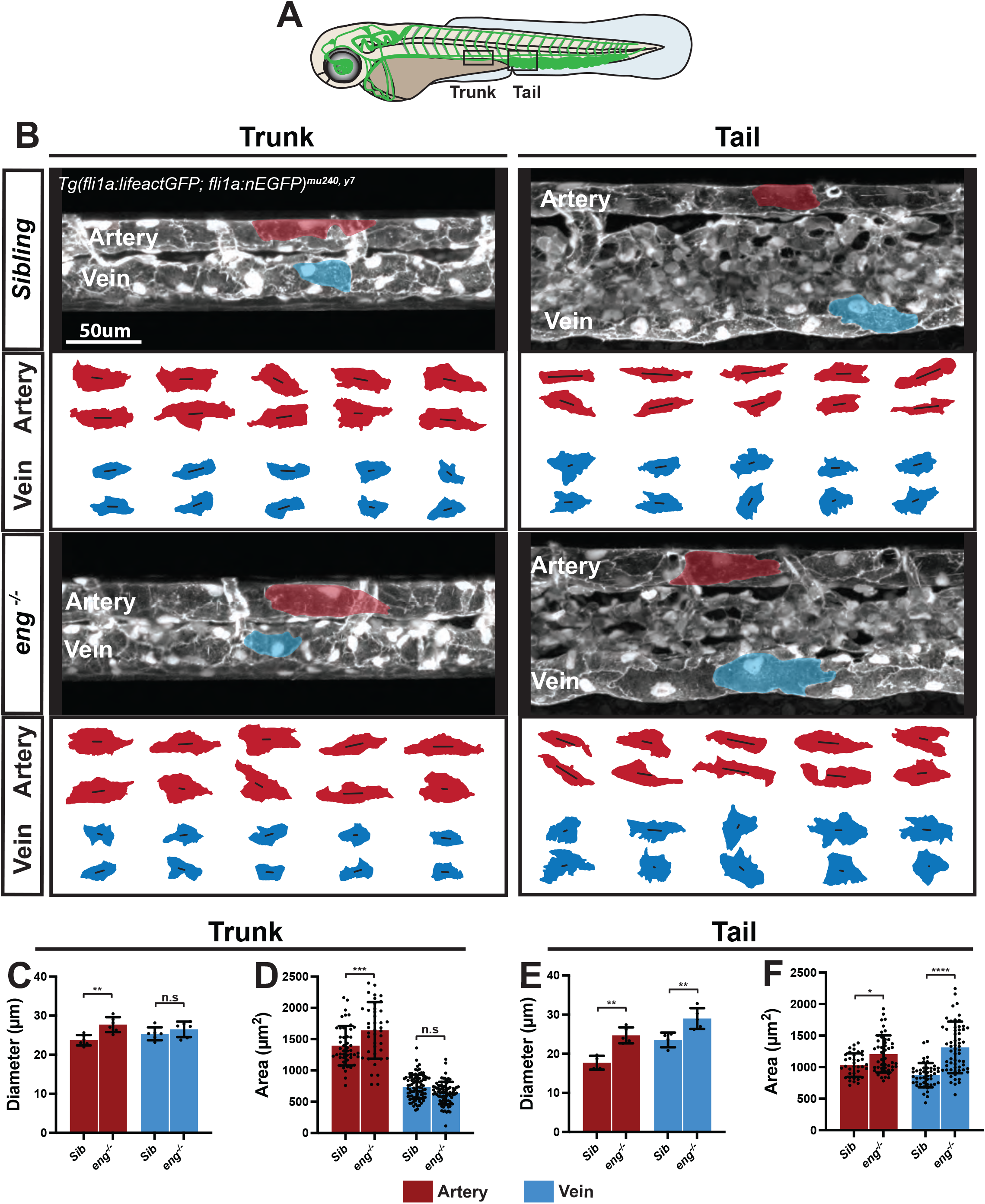
Global loss of endoglin increases EC size in the artery and vein. (A) Schematic of a 3dpf zebrafish embryo with the vasculature highlighted in green. The outlined solid boxes indicate trunk and tail regions. (B) Confocal images of the trunk and tail major artery and vein at 3dpf in eng^-/-^ mutants and siblings. 2D cell outlines shown below images. Length and angle of black line represent elongation and alignment of cells, respectively. (C-F) Quantifications of vessel diameters and EC surface areas in the trunk (C, D) and tail (E, F) of eng^-/-^ mutants and siblings. Data analyzed by one-way ANOVA. n.s, not significant; *P <0.05 **P <0.01, ***P <0.001 ****P <0.0001; error bars indicate s.d.

### Endoglin functions cell autonomously to restrict venous EC size

We previously showed that arterial blood flow is higher in *eng* mutants (Sugden et al., 2017). We therefore hypothesize that the increase in arterial EC size in *eng* mutants might be a secondary effect of increased blood flow. To separate the effects of *eng* loss of function from those of increased flow in a vessel-specific manner, we generated genetic mosaics, in which only a few ECs were mutant in an otherwise WT vasculature. Briefly, we transplanted cells from *eng* mutant or WT donor embryos into WT recipients and determined the morphology of donor ECs in mosaic embryos at 3 dpf **(Fig. 4A)**. We observed that transplanted mutant arterial ECs were not significantly different in size compared to WT arterial ECs. By contrast, transplanted mutant venous ECs were significantly larger than their WT counterparts in both the trunk and the tail **(Fig. 4B-E)**. No difference was observed in elongation and alignment between mutant and WT transplanted ECs **(Suppl. Fig. 4A, B)**. This indicates that expression of Endoglin in veins is cell-autonomously required to restrict venous EC size and that the arterial EC size increase in *eng* mutants is most likely caused by increased flow.

**Figure 4.**
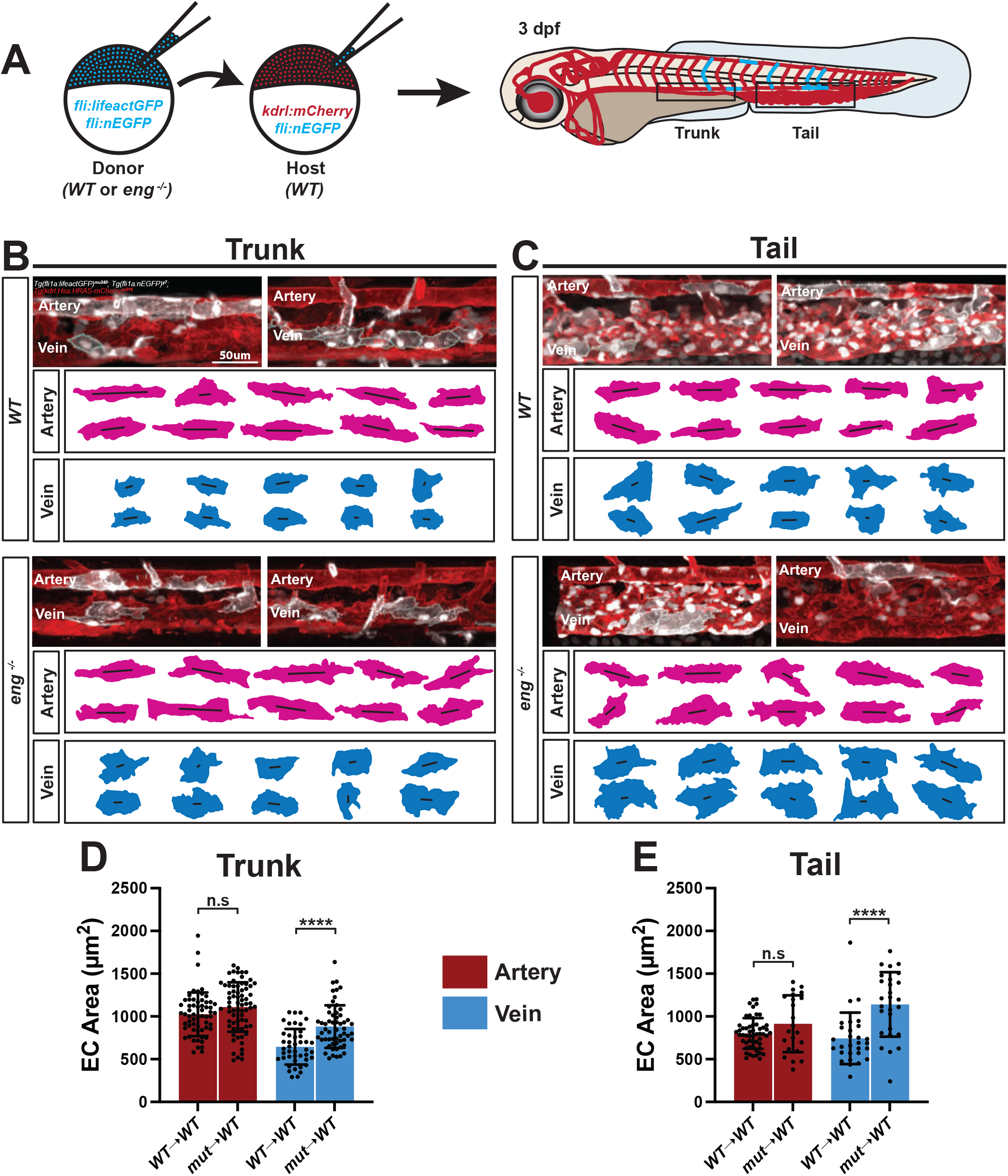
Endoglin regulates venous endothelial cell size in a cell autonomous manner. (A) Schematic of transplantation of WT or *eng*^-/-^ mutant ECs into WT host. Right hand side shows a 3dpf mosaic host zebrafish embryo with the vasculature highlighted in red and transplanted ECs in cyan. The outlined solid boxes indicate the trunk and tail regions analyzed. (B, C) Confocal images of the trunk (B) and tail (C) major artery and vein at 3dpf in WT host embryos transplanted with WT or *eng*^-/-^ mutant ECs. 2D cell

### Alk1 functions cell autonomously to restrict venous EC size despite its predominantly arterial expression

*alk1* mutant zebrafish embryos have a large arteriovenous shunt in the head that reduces blood flow to the axial vessels (Roman et al., 2002). Consequently, we would not be able to separate the effects of reduced flow from those caused by loss of Alk1 on arterial and venous EC sizes. We therefore performed cell transplantations to unravel the primary effects of loss of Alk1 on arterial and venous EC size independently of flow changes **(Fig. 5A)**. We surprisingly discovered that *alk1* mutant transplanted ECs behaved very similarly to *eng* mutant transplanted ECs. Mutant arterial ECs were not significantly different in size compared to WT cells. Conversely, mutant venous ECs were nearly 50% larger in both the trunk and the tail **(Fig. 5B-E)**. Elongation and alignment were comparable between mutant and WT ECs **(Suppl. Fig. 5A, B)**. These observations stand in contrast to the pronounced arterial expression pattern of Alk1 and indicate that low levels of Alk1 expression in the vein might be sufficient to restrict venous EC size, likely through interaction with Endoglin.

**Figure 5.**
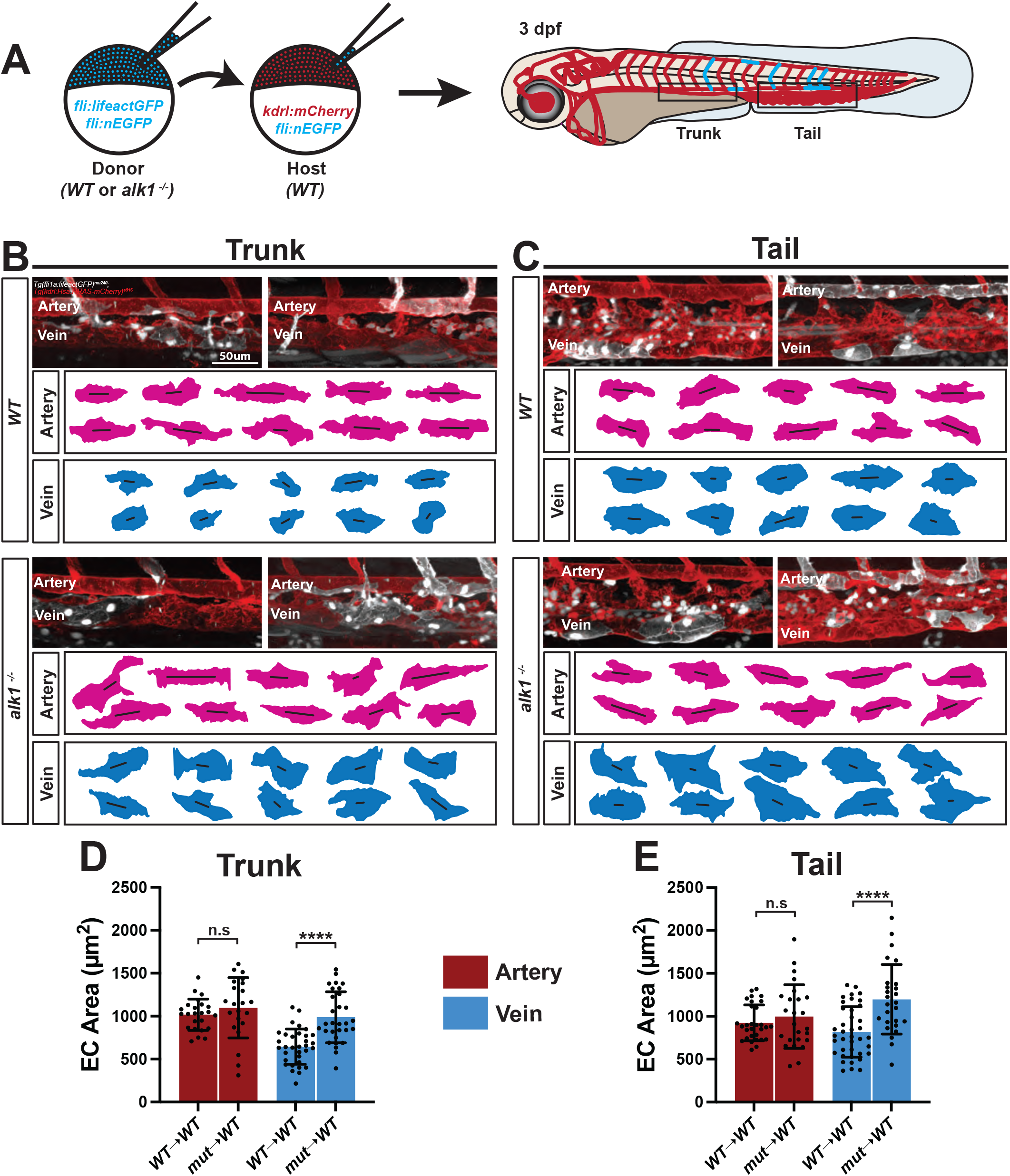
Alk1 regulates venous endothelial cell size despite higher expression in arteries. (A) Schematic of transplantation of WT or *alk1*^-/-^ mutant ECs into WT host. Right hand side shows a 3dpf mosaic host zebrafish embryo with the vasculature highlighted in red and transplanted ECs in cyan. The outlined solid boxes indicate the trunk and tail regions analyzed. (B, C) Confocal images of the trunk (B) and tail (C) major artery and vein at 3dpf in WT host embryos transplanted with WT or *alk1*^-/-^ mutant ECs. 2D cell

### Initiation of vessel dilation in *endoglin* mutants occurs in the tail vein

To further probe our hypothesis that the primary dilation in *eng* mutants is in the vein, secondarily causing arterial diameter expansion through flow increases, we performed a time resolved analysis of the tail arterial and venous diameters to determine which vessel dilates first. We performed timelapse imaging of the tail vessels in WT and *eng* mutant embryos from 50-72 hours post fertilization (hpf) and quantified vessel diameters at hourly intervals. As expected, we observed that while WT vessels shrank in diameter during this time interval, *eng* mutant arteries and veins dilate **(Movies 1-4)**. While quantifying vessel diameters in the WT tail region, we noted a steep drop in diameters along each vessel from anterior to posterior. This introduced a larger range of variability in diameter calculations and prevented an accurate determination of diameter changes over time. To circumvent this issue, we divided the tail region into an anterior and a posterior segment, each consisting of around 6 somitic segments **(Fig. 6A)**.

**Figure 6.**
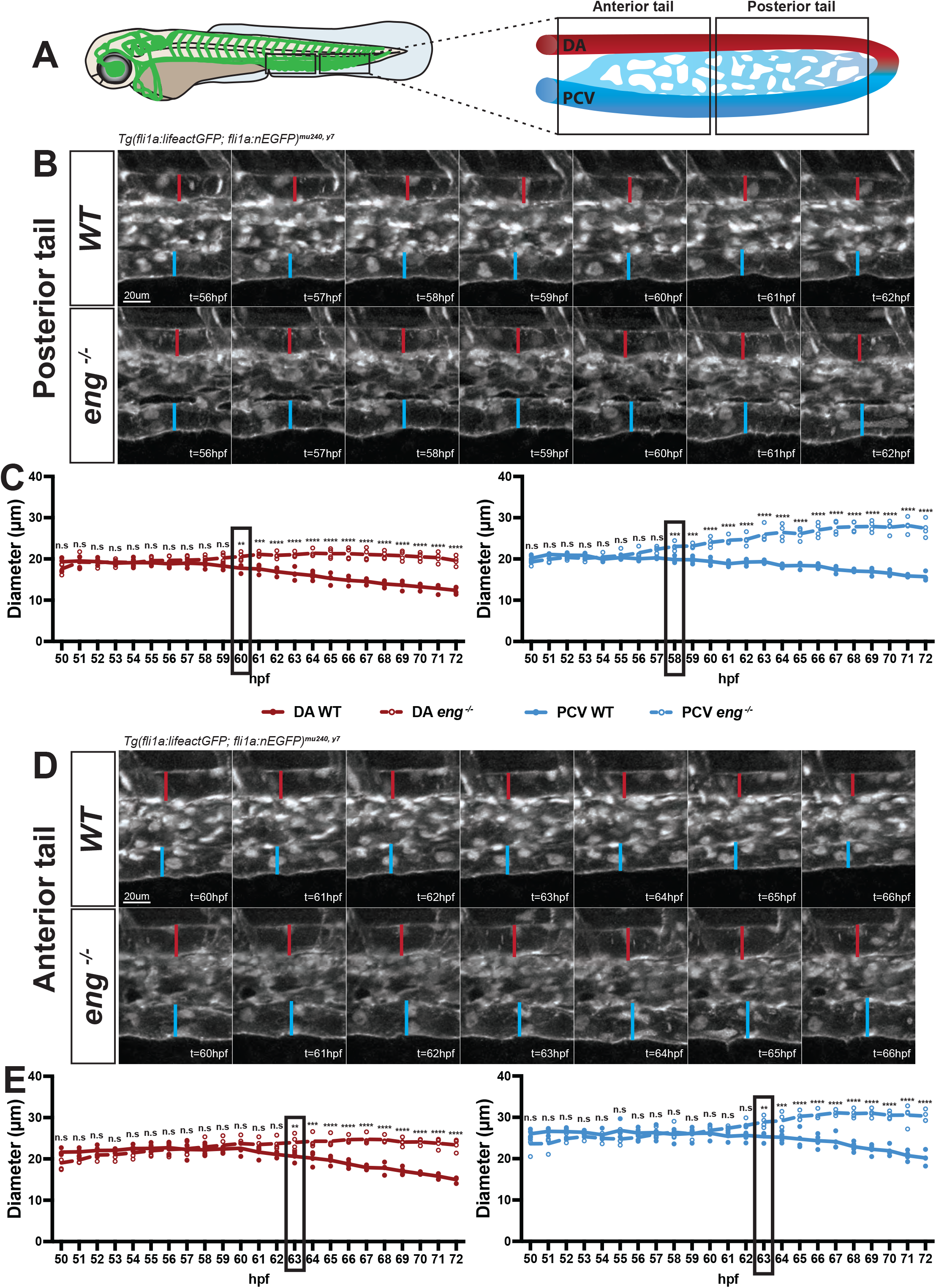
Initiation of vessel dilation in endoglin mutants occurs in the tail vein. (A) Schematic of a 3dpf zebrafish embryo with the vasculature highlighted in green. The black outlined boxes indicate anterior and posterior sections of the tail vasculature with magnified views on the right. (B, D) Timelapse images of WT and *eng*^-/-^ mutant artery and vein in the posterior (B) and anterior (D) tail regions. Vertical red and blue lines indicate diameter of artery and vein, respectively. (C, E) Quantifications of artery and vein diameters in the posterior (C) and anterior (E) tail region of WT and *eng*^-/-^ mutants. Diameters quantified per hour from 50-72hpf timelapse movies. Black outlined boxes represent the timepoints where *eng*^-/-^ mutant and WT diameters start diverging significantly. Data analyzed across genotypes by two-way ANOVA. n.s, not significant; *P <0.05, **P <0.01, ***P <0.001, ****P <0.0001; error bars indicate s.d.

We noticed a larger increase in *eng* mutant venous diameter compared to arterial in both tail segments. We also observed a difference in the timepoint when mutant diameters diverged from WT for the artery versus the vein. The first increase in *eng* mutant vessel diameter was initiated in the posterior tail vein around 58 hpf, followed by the posterior tail artery around 60 hpf **(Fig. 6B, C; Suppl. Fig. 6A)**. Subsequently, the mutant anterior tail artery and vein showed a divergence from WT around 63 hpf **(Fig. 6D, E; Suppl. Fig. 6A)**. This data, combined with our earlier observation that Endoglin cell autonomously regulates venous EC size, provides evidence that the earliest effect of loss of Endoglin is an expansion of venous diameter through an increase in venous EC size, and that venous dilation initiates in the posterior region of the embryo that actively undergoes angiogenesis. The increase in arterial diameter and EC size is a secondary effect, most likely caused by aberrant increases in blood flow brought about by increased venous diameter.

## Discussion

Despite the importance of maintaining vascular hierarchy, the underlying cellular and molecular mechanisms that spatiotemporally establish blood vessel diameters have remained unclear. Mural cells – with their plethora of cell types in differing vascular localizations – are known to regulate blood vessel dilation and constriction in response to shear stress (Siekmann, 2023). However, mural cells do not mature until later in development, and in the zebrafish axial vasculature this occurs at around 3 dpf (Ando et al., 2021, 2019). Hence, murals cells are unlikely to contribute to vessel diameter regulation during earlier developmental stages. We thus set out to explore the mechanisms regulating vessel diameters during early embryonic development (1-3 dpf in zebrafish, period of vessel formation and initial remodelling). Since ECs are the only cell type lining blood vessels during this time, we focused on the contribution of ECs to this process.

Our findings reveal the importance of EC size in spatiotemporally controlling vessel calibre during early development. Moreover, our study underscores the requirement for BMP pathway components – Endoglin and Alk1 – in specifically controlling venous diameters to prevent AVMs during development. A recent studies in mice have also shown that loss of Eng or Alk1 from veins and/or capillaries is the underlying cause for AVMs (Jin et al., 2017; Park et al., 2021; Singh et al., 2020). Specifically, a pan-endothelial or venous/capillary-specific loss of Eng expression leads to a similar incidence and frequency of AVMs in the postnatal mouse retina (Singh et al., 2020). Moreover, AVMs arising from pan-endothelial loss of Eng cannot be rescued by arterial ECs overexpressing human ENG (hENG) and a complete rescue requires hENG overexpression in veins and capillaries as well (Jin et al., 2017). Furthermore, deletion of Alk1 specifically from veins and capillaries was sufficient to induce AVMs in the postnatal retina due to impaired EC polarization against flow (Park et al., 2021). These studies highlight the requirement for Eng and Alk1 in veins and capillaries for proper EC polarization and migration against flow to prevent AVMs. Our study adds to this knowledge by taking advantage of the zebrafish model system to uncover a novel and cell-autonomous requirement for Eng and Alk1 in restricting venous EC size during early development to prevent AVMs. Our study also elaborates on the real-time interaction between hemodynamic forces and EC size to precipitate AVMs in a spatiotemporal manner.

It is interesting to note is that mouse models of *eng* or *alk1* LOF display increased EC proliferation or defective EC migration as the primary mechanism for AVMs (Baeyens et al., 2016; Hwan Kim et al., 2020; Jin et al., 2017; Mahmoud et al., 2010; Ola et al., 2016; Park et al., 2021; Tual-Chalot et al., 2014). However, the results of our current study in zebrafish suggest that the earliest defect is an increase in venous EC size without a change in EC number. These seemingly contradictory results in the two model systems might arise due to one or more of the following reasons: differences in developmental stages at which the phenotypes were observed, presence of tissue-specific phenotypes, or exposure of ECs to different environmental cues in the two model systems. Our previous study offers some clue to understanding these differences (Sugden et al., 2017). There, we observed higher venous EC numbers in the adult fin vasculature of *eng* mutants, and not in the embryonic tail that the fin arises from. This points toward a role for EC number in AVM pathogenesis later in development, with EC size being the chief contributor to vessel dilation during early development.

Another intriguing observation about AVMs is that they can be mosaic for the genetic mutation. Recent studies on mouse models of HHT1 involving *eng* LOF or of CCM involving *ccm3* LOF have noted that AVMs consist of a combination of mutant and WT cells (Jin et al., 2017; Malinverno et al., 2019). This raises several questions: Is recruitment of WT cells to AVMs a passive process or are they actively enlisted by mutant cells, and if so, by what mechanisms? Do WT cells change their behaviours and/or functions within affected vessel segments? How many mutant cells are sufficient to precipitate an AVM? Further experimental insights into these questions might help explain the heterogeneity of phenotypes observed in HHT patients.

Previous studies have linked AVM formation in *eng* or *alk1* LOF mouse models to an upregulation of the PI3 kinase (PI3K) pathway (Jin et al., 2017; Ola et al., 2016). Interestingly, despite its well-known role in the regulation of EC proliferation and migration (Kobialka and Graupera, 2019), the effect of PI3K pathway on EC size remains uncertain (Angulo-Urarte et al., 2018; Crackower et al., 2002; di Blasio et al., 2018; García et al., 2006; Neufeld, 2003). It is, however, well established that the downstream mTOR axis of PI3K pathway promotes cell growth through protein and ribosomal synthesis (Liu and Sabatini, 2020; Wang and Proud, 2006). Thus, further investigation into the contribution of the PI3K-mTOR pathway to EC size regulation may reveal novel avenues to understanding the mechanisms regulating EC size downstream of the BMP pathway.

The shear stress set point theory has been widely proposed as a mechanism for vessels to maintain homeostatic shear stress levels in response to changes in flow. However, an *in vivo* shear stress set point has only been studied and noted to exist in arteries. The existence of an *in vivo* set point for veins has remained an open and surprisingly unasked question. We show in our study that the major axial vein displays context-dependent responses to flow, with the anterior, more mature vein maintaining its diameters and EC sizes; while the posterior, more angiogenic vein responds to both increases and decreases in flow, albeit to a lower extent than the artery. Our findings indicate the presence of a spatiotemporally restricted and less defined *in vivo* shear stress set point for veins – a hypothesis that warrants further investigation.

Thus, based on findings from our current study, we propose a model wherein expression of Eng and Alk1 in venous ECs signals as an EC size brake and limits the response of veins to blood flow. Arterial ECs, on the other hand, respond to flow by increasing their size. Consequently, the continual increase in blood flow during early development only causes arterial expansion with no changes in venous diameter, allowing systemic pressure to be maintained and therefore smaller vessels to be perfused **(Fig. 7A)**. Loss of Endoglin or Alk1 causes veins to behave like arteries by triggering a dramatic increase in venous EC size and thus diameter in response to flow. The resulting influx of blood flow leads to an amplified response within arterial ECs, further increasing their size and thus arterial diameter. The combined expansion of both arterial and venous vessels brings in excess flow that triggers further vessel expansion. This cycle ultimately precipitates an arteriovenous shunt **(Fig. 7B)**.

**Figure 7.**
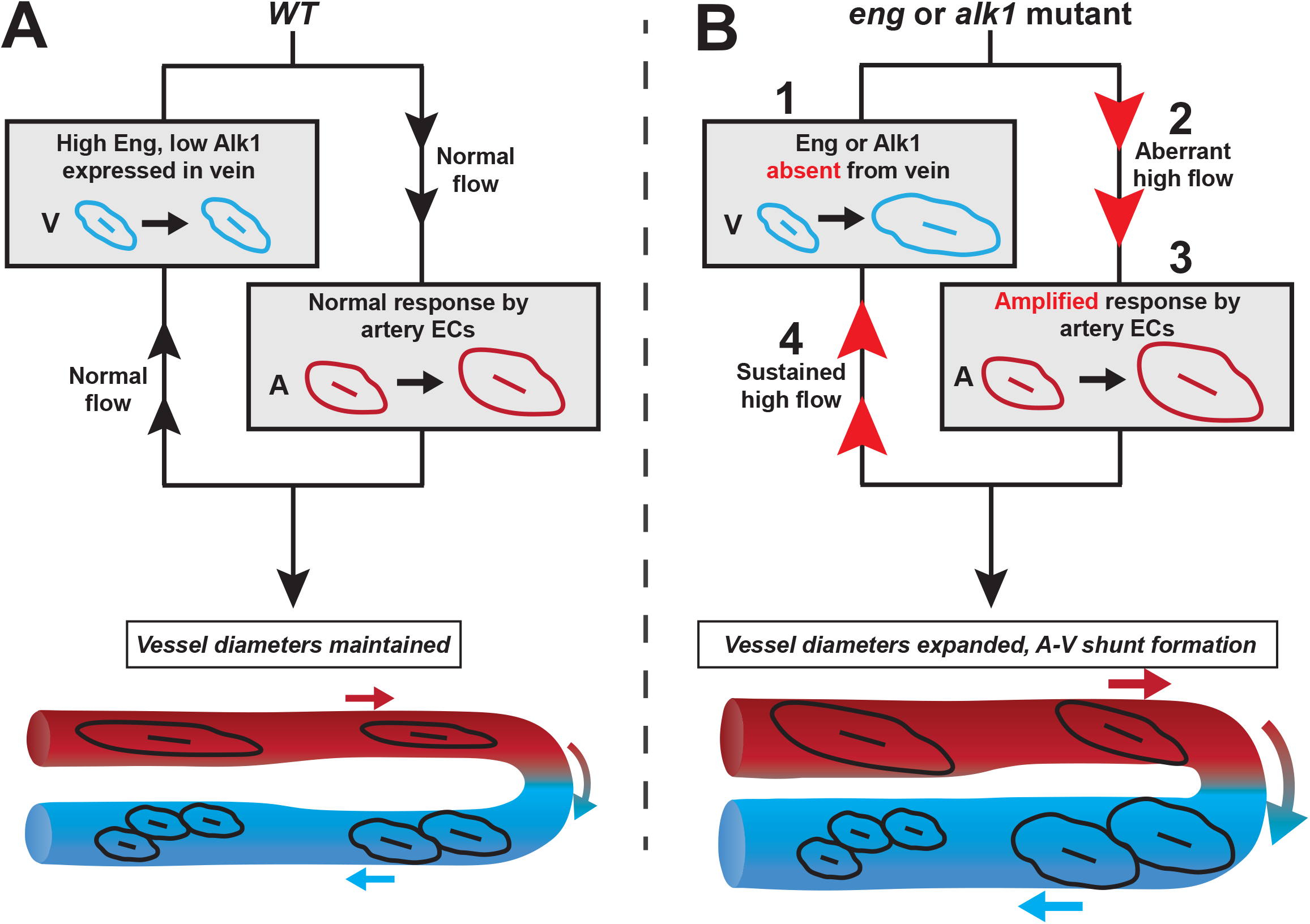
Proposed model for role of Endoglin and Alk1 in regulating venous EC size. (A) In the WT vasculature, high levels of Endoglin and low levels of Alk1 in venous ECs signal as an EC size brake and prevent the vein from expanding in response to flow. The arterial ECs, on the other hand, respond normally to flow by increasing their size. The resultant increase in arterial diameter but not venous allows flow to be distributed to smaller vessels. (B) Loss of Endoglin or Alk1 from the vein allows venous ECs to increase their size in response to flow (1). The resulting increase in venous diameter lets in more flow (2) and causes arterial ECs to secondarily expand their size as an amplified response to higher-than-normal flow (3). The combined dilation of both the artery and vein acts as a shunt (4) and prevents blood flow from entering smaller vessels, leading to an arteriovenous malformation. leading to an arteriovenous malformation.

We found it intriguing that despite its higher expression in the artery, Alk1 is required for EC size control in the vein. One possible explanation for this finding involves the role of Endoglin, a coreceptor, in potentiating signalling through Alk1 (Blanco et al., 2005; Lebrin and Mummery, 2008). We speculate that low levels of Alk1 alone might be able to initiate pathway activation but require Eng to increase signalling to prevent EC size increases. Since this condition is satisfied in the vein, the primary function of Alk1 and Eng seems to be in this vessel. Conversely, despite higher Alk1 in the artery, it is possible that there is not enough Eng to sustain pathway activation and consequently restrict arterial EC size. Our study thus emphasizes the emerging role of veins as a crucial player in the precipitation of vascular malformations and could potentially inform the development of novel targeted therapies for the treatment of HHT.

## Methods

### Zebrafish husbandry and strains

All research protocols involving zebrafish were approved by the University of Pennsylvania Institutional Animal Care and Use Committee. Zebrafish embryos were maintained in 1X E3 medium under recommended animal husbandry conditions (Westerfield, 2000) and in accordance with institutional and national regulatory standards. Transgenic lines and mutants used were *Tg(fli1a:lifeactGFP)^mu^*^240^, *Tg(fli1a:nEGFP)^y^*^7^, *Tg(kdrl:Hsa.HRAS-mCherry)^s^*^916^, *eng^mu130^*, and *alk1^y6^*. References for all zebrafish lines can be obtained on http://zfin.org.

### Live imaging and confocal microscopy

1-3 dpf embryos were mounted on a glass-bottomed dish using 1% low-melting-point agarose containing 168 mg/l tricaine and 0.003% phenylthiourea (to prevent pigment formation). For time lapse imaging, a heated microscope stage was used to maintain a constant temperature of 28.5°C. Confocal *z*-stacks were acquired on the SP8-inverted (Leica Microsystems) scanning confocal microscope.

### Tricaine treatments

Embryos were dechorionated and treated with 0.5mg/ml tricaine methanesulfonate (Millipore Sigma) in 1X E3 containing 0.003% phenylthiourea. Treatments were performed as staggered experiments with each group of embryos treated for 2h intervals at 1, 1.5 or 2dpf, and imaged immediately after at the respective time points, i.e., 26, 38 or 50hpf.

### Image processing

Imaris software (Oxford Instruments) was used for image analysis, including maximum intensity projections, timelapse movies, cell shape tracings, and measurement of vessel diameters, cell numbers and cell proliferation. Adobe Illustrator software was used to assemble figures and create schematics.

### Vessel diameter, cell number and cell shape analysis

To calculate vessel diameters, measurements were taken at the midway point between ISVs no. 7-15 for trunk and ISVs no. 16-24 for tail. The mean was used as the average diameter per embryo per trunk or tail region. Nuclei counts were obtained for 430µm length of each vessel between ISVs no. 7-15 for trunk and ISVs no. 16-24 for tail.

Cell shape analysis was performed as described previously (Sugden et al., 2017). Briefly, *Tg(fli1a:lifeactGFP; fli1a:nEGFP)^mu^*^240,*y*7^ double transgenic embryos were imaged with a 40X air or 60X water immersion objective. Imaris software was used to manually trace cell outlines with an average of 100 measurement points per cell. Using a custom MATLAB script, these points were then unrolled onto a two-dimensional surface, and an ellipse or a hyperbola was fitted through the measurement points. Cell area, elongation and alignment were then calculated from the two-dimensional projection.

### Proliferation rate

To quantify cell proliferation rate, time lapse movies were analyzed for cells that displayed nuclear dissociation and subsequent cell division. Proliferation events were counted for each vessel between 1-1.5, 1.5-2 and 2-3dpf and normalized to the total cell number (per 430µm vessel length) to obtain rate of proliferation.

### Blastomere transplantations

Cell transplantations were performed as described previously (Siekmann and Lawson, 2007). Wildtype, *eng^mu130^* mutant or *alk1^y6^* mutant embryos with a *Tg(fli1a:lifeactGFP; fli1a:nEGFP)^mu^*^240,*y*7^ background were used as donors, while recipients were wildtype *Tg(kdrl:Hsa.HRAS-mCherry; fli1a:nEGFP)^s^*^916,y7^.

### Whole mount *in situ* hybridization

Primer sequences for amplifying *alk1* were kindly provided by the lab of Beth Roman (University of Pittsburgh, Pittsburgh). *eng* probe was generated as described previously (Sugden et al., 2017). Whole-mount *in situ* hybridization of zebrafish embryos was performed as described previously (Hauptmann and Gerster, 1994). ‘*n*/*n*’ reports the ‘number of embryos with staining pattern in image’/‘total embryos from 2 experiments’.

### Statistical analysis

All data were analyzed using GraphPad Prism and graphs were plotted with mean standard deviation (s.d.). *P*<0.05 was considered statistically significant.

## Code availability

All code used in this study is available from the authors upon request.

## Notes

### Competing Interest Statement

The authors have declared no competing interest.

