## Supplemental Figures for "Alk1/Endoglin dependent increases in vein endothelial cell sizes precipitate arteriovenous malformations"

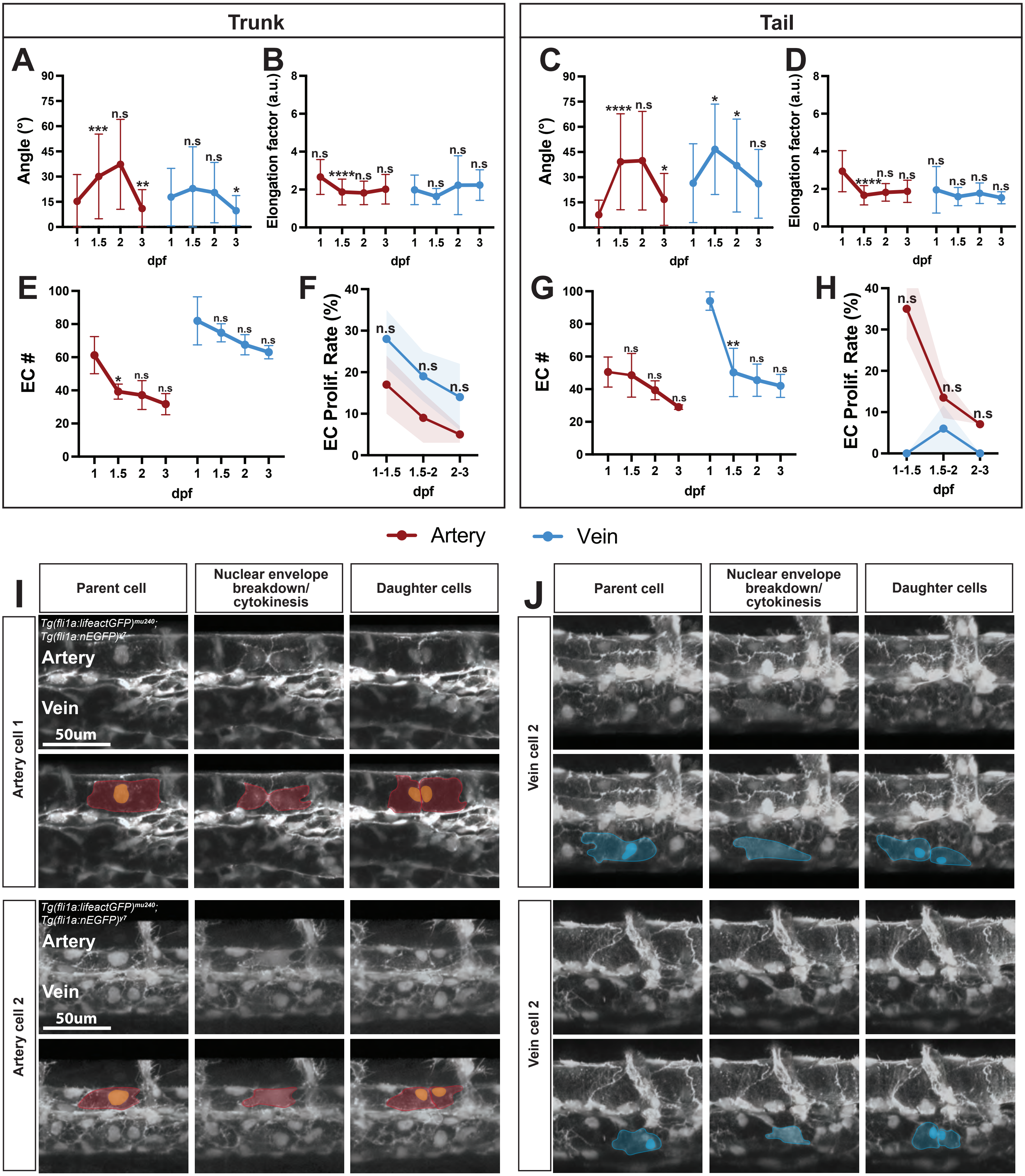

Trunk
Tail
A
B
C
D
E
F
G
H
Angle (°)
Elongation factor (a.u.)
EC #
EC Prolif. Rate (%)
dpf
n.s
Artery
Vein
I
J
Parent cell
Nuclear envelope breakdown/ cytokinesis
Daughter cells
Tg(fli1a:lifeactGFP)mu240; Tg(fli1a:nEGFP)y7
Artery
Vein
50um
Artery cell 1
Artery cell 2
Vein cell 2

**Suppl. Figure 1. Arterial and venous ECs have similar proliferation rates during early development**
(A-D) Quantifications of EC alignment, elongation, number and proliferation rate in the trunk (A, B, E, F) and tail (C, D, G, H) major artery and vein. Data analyzed across developmental stages for (A-E, G) and across vessel types for (F, H) by two-way ANOVA. n.s, not significant; *P <0.05, **P <0.01, ***P <0.001, ****P <0.0001; error bars and shaded red and blue regions indicate s.d. (I, J) Representative snapshots of EC proliferation events (before, during and after cell division) from timelapse videos of the trunk artery and vein between 1.5-2dpf. Arterial ECs pseudo-colored red, venous ECs blue.

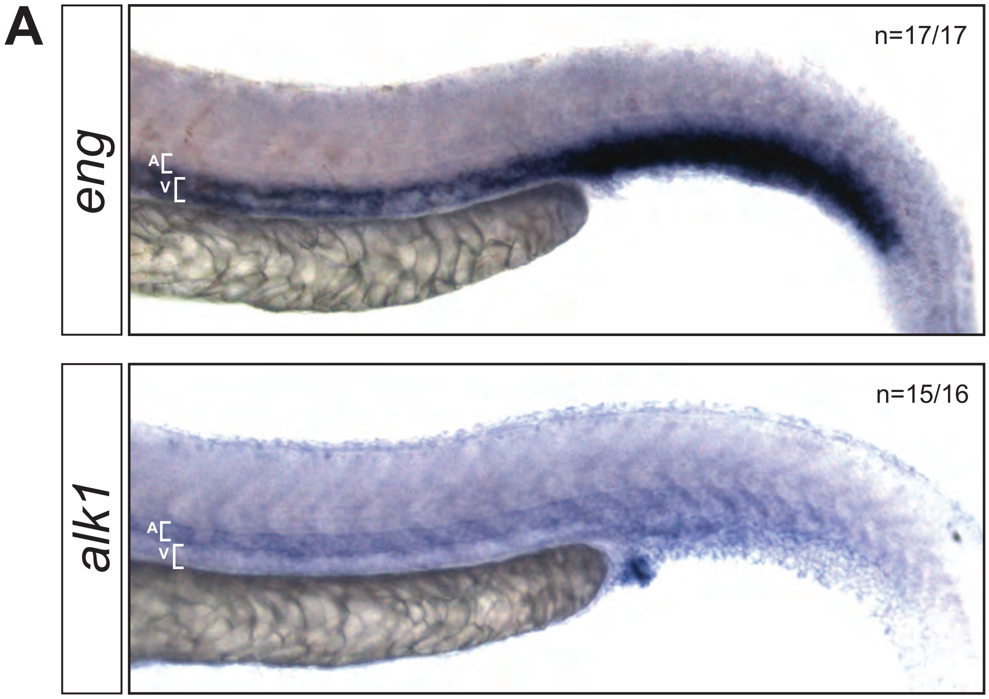


**Suppl. Figure 2. Endoglin is expressed predominantly in veins and alk1 in arteries**
(A) Whole mount *in situ* hybridization for *endoglin* and *alk1* in 1.5dpf *WT* embryos. A = artery, V = vein.

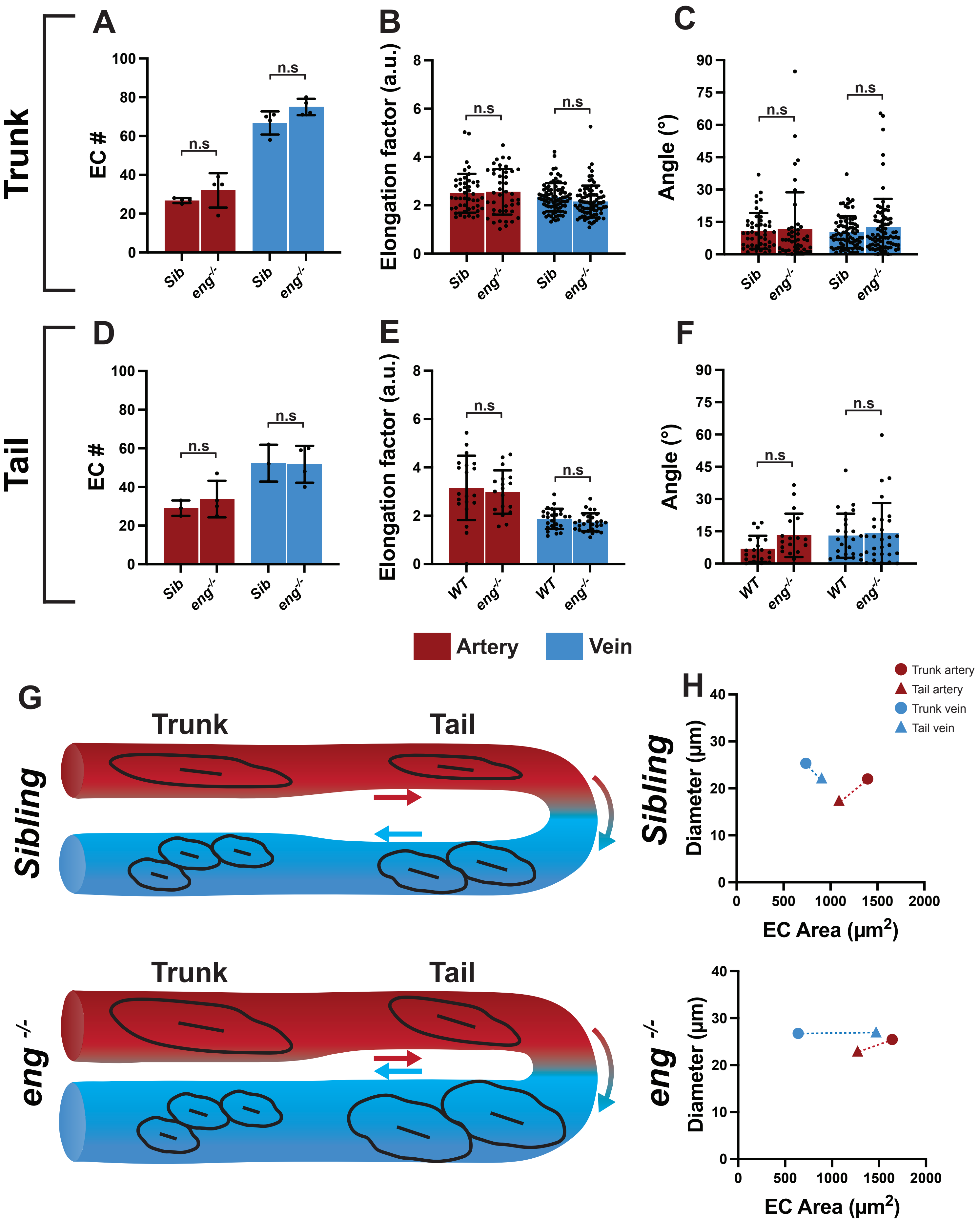

Trunk
Tail
A
B
C
D
E
F
EC #
Elongation factor (a.u.)
Angle (°)
n.s
Sib
eng-/-
WT
Artery
Vein
G
Trunk
Tail
Sibling
eng -/-
H
Trunk artery
Tail artery
Trunk vein
Tail vein
Diameter (µm)
EC Area (µm2)

**Suppl. Figure 3.** (A-F) Quantifications of EC number, elongation and alignment in the major artery and vein of *eng*$^{-/-}$ mutant embryos and siblings in the trunk (A-C) and tail (D-F) regions at 3dpf. Data analyzed by one-way ANOVA, n.s, not significant; *P <0.05, **P <0.01, ***P <0.001, ****P <0.0001; error bars indicate s.d. (G) Schematic of EC morphologies in the major artery and vein of *eng*$^{-/-}$ mutant and sibling embryos at 3dpf. (H) Diameter-to-EC area correlation graphs for *eng*$^{-/-}$ mutant and sibling embryos. Circle = trunk, triangle = tail, blue = vein, red = artery.

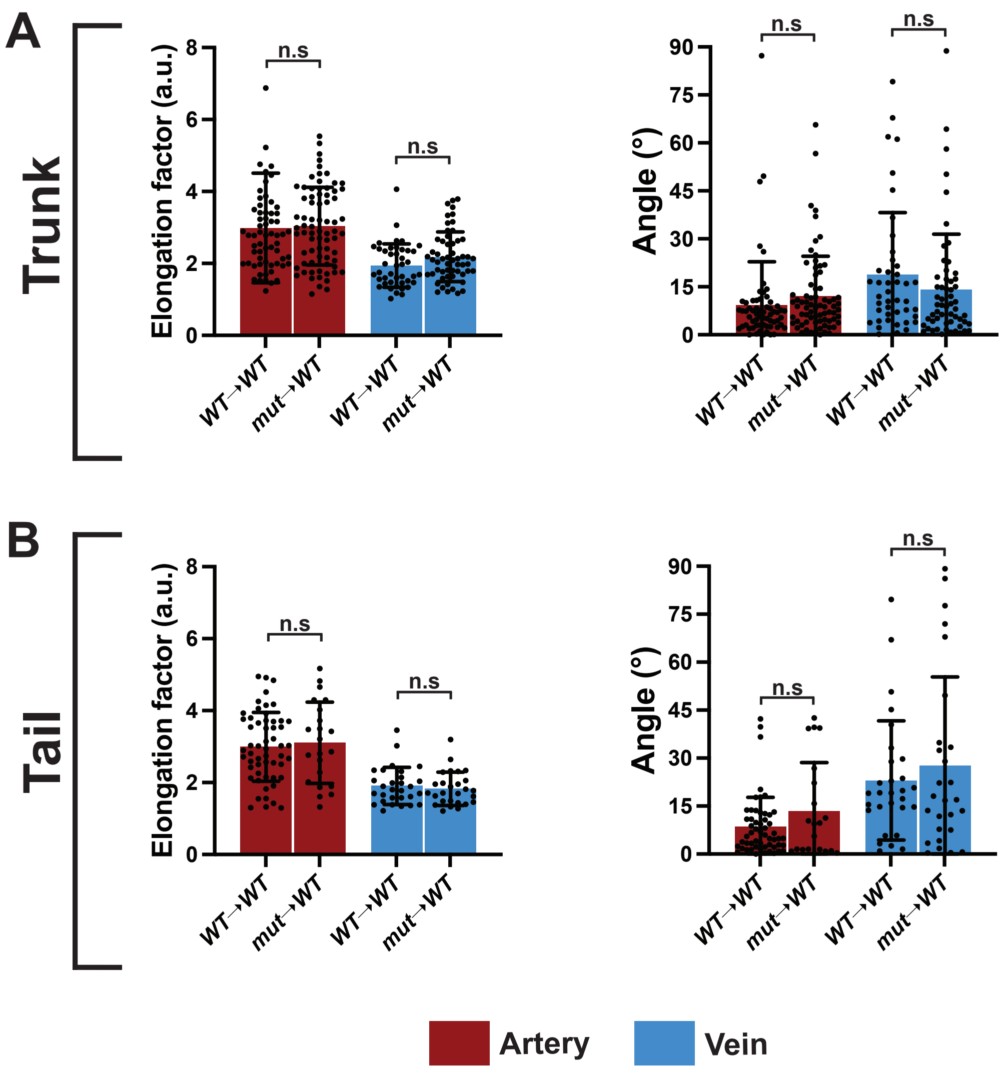


**Suppl. Figure 4.** (A, B) Quantifications of elongation and alignment of ECs transplanted from *WT* or *eng*$^{-/-}$ mutant embryos into the trunk (A) and tail (B) of *WT* host. Data analyzed by one-way ANOVA. n.s, not significant; error bars indicate s.d.

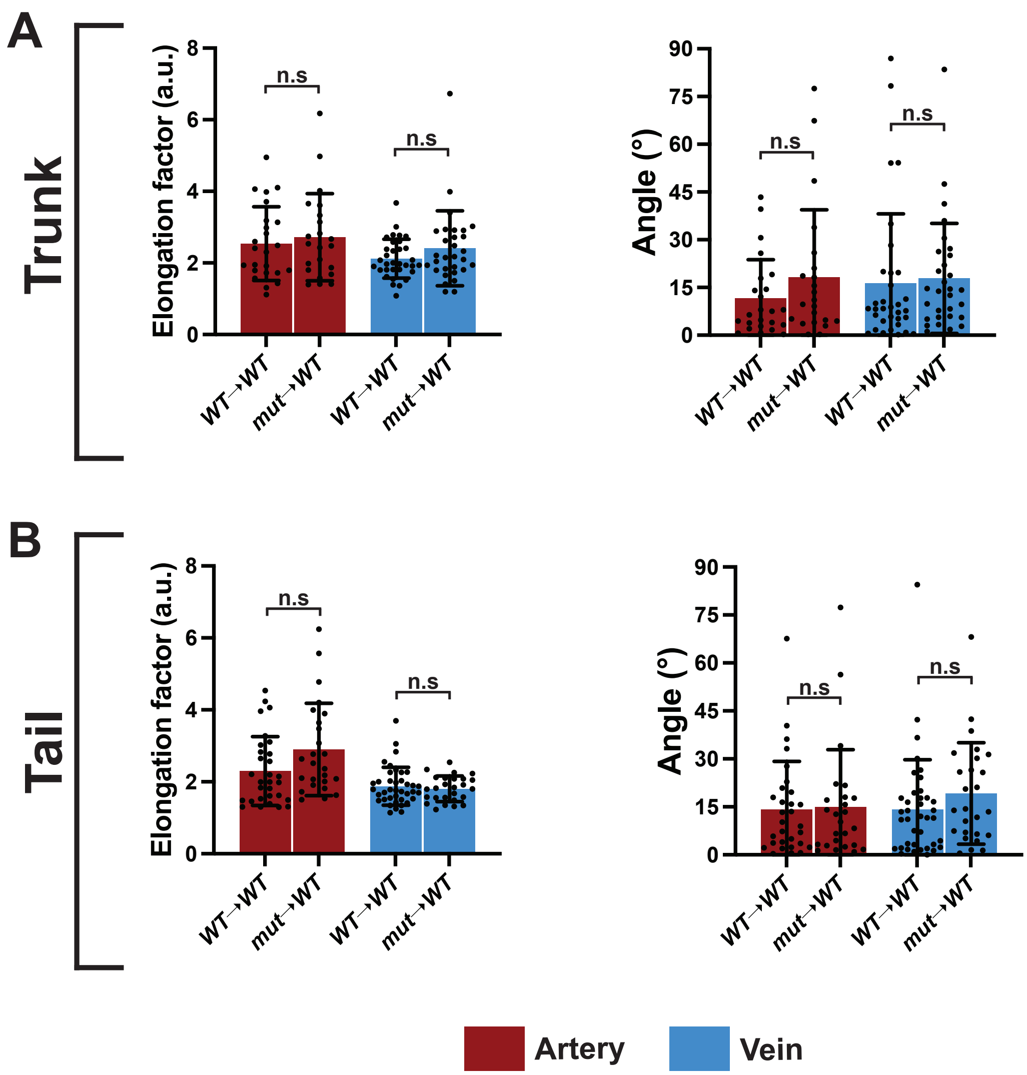


**Suppl. Figure 5.** (A, B) Quantifications of elongation and alignment of ECs transplanted from *WT* or *alk1*$^{-/-}$ mutant embryos into the trunk (A) and tail (B) of *WT* host. Data analyzed by one-way ANOVA. n.s, not significant; error bars indicate s.d.

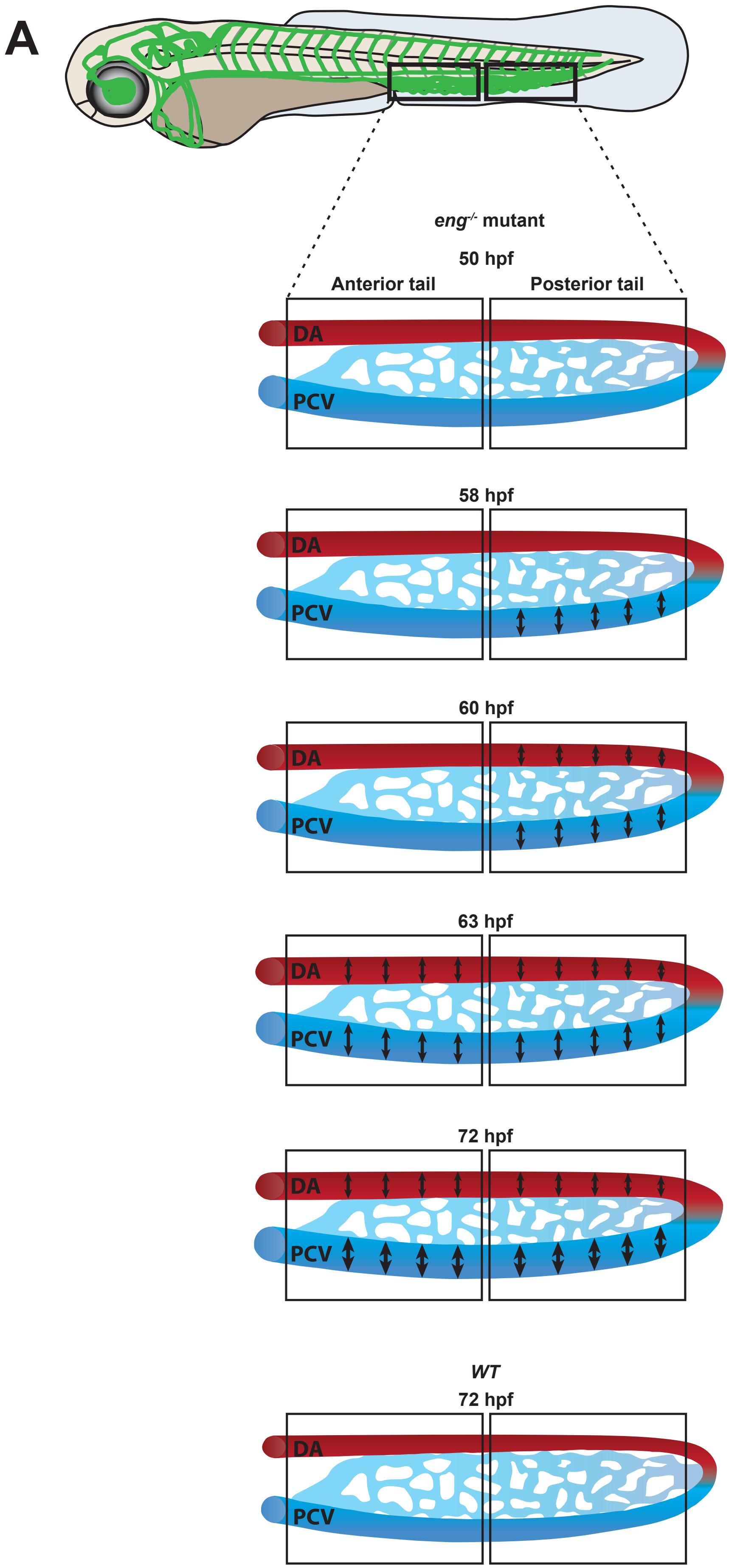
A
eng-/- mutant
50 hpf
Anterior tail
Posterior tail
DA
PCV
58 hpf
DA
PCV
60 hpf
DA
PCV
63 hpf
DA
PCV
72 hpf
DA
PCV
WT
72 hpf
DA
PCV

**Suppl. Figure 6. Schematic of initiation of vessel dilation in endoglin mutants**

(A) Schematic of a 3dpf zebrafish embryo with the vasculature highlighted in green. The black outlined boxes indicate anterior and posterior sections of the tail vasculature. Magnified schematics of the tail vasculature below depict the sequence of events during shunt formation in *eng*$^{-/-}$ mutants from 50-72hpf.
